# Machine-learning-based automated Schlemm’s canal volumetric segmentation for optical coherence tomography

**DOI:** 10.1101/2025.09.08.674998

**Authors:** Raymond Fang, Fengyuanshan Xu, Zihang Yan, Cheng Sun, Tsutomu Kume, Alex S. Huang, Hao F. Zhang

## Abstract

Volumetric segmentation of Schlemm’s Canal (SC) in optical coherence tomography (OCT) is time-consuming, creating a barrier to experiments studying glaucoma and the anatomy of the trabecular outflow pathways *in vivo*. To this end, we developed an automated segmentation tool, Schlemm’s Canal-Localization and Semantic Segmentation (SC-LSS), for the volumetric segmentation of SC in *in vivo* mice eyes from visible-light OCT (vis-OCT). SC-LSS first localizes the boundaries of SC and subsequently determines the boundaries of SC within the localized region. We used 324 B-scans from 16 mouse eyes for training, validation, and testing the model, and 203 additional B-scans from 16 mouse eyes to evaluate the model’s accuracy. We found that the Dice coefficient between segmentations generated by SC-LSS and manual expert graders was 0.70 ± 0.20 and that the Dice coefficient between two expert graders was 0.73 ± 0.18 (p = 0.10). Furthermore, SC-LSS captured decreases in SC size with increasing intraocular pressure, yielding a 51.5% decrease in SC size at 20 mmHg compared to 5 mmHg. SC-LSS also identified a 20.1% increase in SC size following the administration of pilocarpine. We anticipate that SC-LSS will accelerate studies on factors regulating the trabecular outflow pathways and their role in glaucoma development and management.

## 1. Introduction

Glaucoma, a leading cause of irreversible blindness worldwide, is an optic neuropathy characterized by damage to retinal nerve ganglion cells^1, 2^. Elevated intraocular pressure (IOP) is the primary modifiable risk factor for glaucoma, and lowering IOP is the focus of existing medical and surgical treatments for glaucoma^3^. Physiologically, IOP is regulated by the production and outflow of aqueous humor^4^. The majority of outflow occurs through the trabecular outflow pathway, where aqueous humor drains from the anterior chamber of the eye, through the trabecular meshwork (TM) and Schlemm’s canal (SC), and eventually through episcleral and aqueous veins^5, 6^. Thus, visualizing the anatomy of the outflow pathways *in vivo* may provide insight into IOP regulation and guide clinical glaucoma management^7, 8^.

Due to its small size, positioning, and geometry, *in vivo* imaging of these pathways has historically been challenging. Approximating SC as an ellipse, the length of its minor axis in both mice and humans is often less than ten micrometers^9, 10^. In humans, the major axis varies between 150 µm and 350 µm ^11^, while in the mouse, it varies between 50 and 200 µm^12^. Additionally, SC is located deep within the light-scattering sclera, making it challenging to image SC optically. The outflow pathway extends 360 degrees around the circumference of the eye. Different regions around the circumference of the eye have varying outflow resistance and anatomy^13–16^. Moreover, SC size varies around the circumference of the eye, making volumetric imaging particularly advantageous for representative imaging of these pathways^10^. Optical coherence tomography (OCT) has emerged as a technology capable of performing volumetric imaging with micron-resolution^17, 18^, with several groups having utilized it for *in vivo* SC imaging^19^. We recently developed a visible-light OCT (vis-OCT) for volumetric imaging of the trabecular outflow pathways in mice, with an axial resolution of 1.3 µm and a lateral resolution of 9.4 µm in tissue^9^.

Recently, there has been an increasing focus on *in vivo* imaging of SC in both human and mouse eyes. Human imaging has focused on the relationship between SC size and glaucoma, with investigators revealing a smaller SC in eyes with glaucoma^20, 21^. Additionally, phase-sensitive OCT has been utilized to investigate the deformations of the TM during the cardiac cycle and to evaluate the biomechanical properties of the TM^22^. Since mice and humans share comparable anatomy and physiology of the trabecular outflow pathways^23, 24^, numerous studies have focused on SC imaging in mice. Several investigations have examined the change in SC morphology with IOP modulation^25, 26^, which is related to the stiffness of the TM^27^. Additional evaluation has examined the influence of various drugs, including pilocarpine, rho-kinase inhibitors, and dexamethasone on the structure of SC^27–29^. Finally, other studies have focused on 360-degree imaging of the trabecular outflow pathway, exploring segmental anatomical differences of SC anatomy around the eye^9, 10, 30^. Hence, automated SC segmentation methods would greatly expedite the rate of preclinical studies aimed at evaluating SC *in vivo*.

Numerous challenges, including the variable anatomy of SC, exist in segmenting SC from OCT images. Foremost, SC shares imaging features with other structures within the eye, including the episcleral and aqueous veins, as they are all hyporeflective lumens in OCT images. Additionally, SC is small relative to the other structures within the anterior segment of the eye, including the cornea and sclera, as noted above. Regarding SC segmentation in humans, region growing, hidden Markov Chain segmentation, and level set methods have been applied to semi-automate SC segmentation^31–33^. However, these methods often required manual intervention and were evaluated on small datasets. For mice, Choy *et al*. developed a deep learning method for segmenting SC^34^. However, their study segmented individual B-scans rather than volumes, and the images consisted of a much smaller field-of-view (FOV). We sought to leverage advanced deep learning to achieve volumetric segmentation of SC.

In this work, we introduce the Schlemm’s Canal-Localization and Semantic Segmentation (SC-LSS), a deep learning model for volumetric segmentation of SC in vis-OCT volumes in mouse eyes. SC-LSS consists of two sequential networks: the first one identifies the boundaries of the SC, and the second one generates the semantic segmentation masks of the SC. We demonstrated that SC-LSS yields comparable segmentations to three expert graders. Furthermore, SC-LSS detected changes in SC morphology in response to IOP modulation and pilocarpine administration.

## 2. Method

### 2.1 Segmentation pipeline overview

For localization and segmentation of SC (Fig. 1a), we adopted a two-stage pipeline comprised of object localization (Fig. 1b) by the You Only Look Once, version 8 (YOLOv8) model^35^, a real-time object detection model that localizes target objects within images by predicting bounding boxes, followed by prompt-based semantic segmentation (Fig. 1c) using the Segment Anything Model (SAM), a segmentation model that generates pixel-wise masks of target objects based on input prompts^36^.

**Figure 1.**
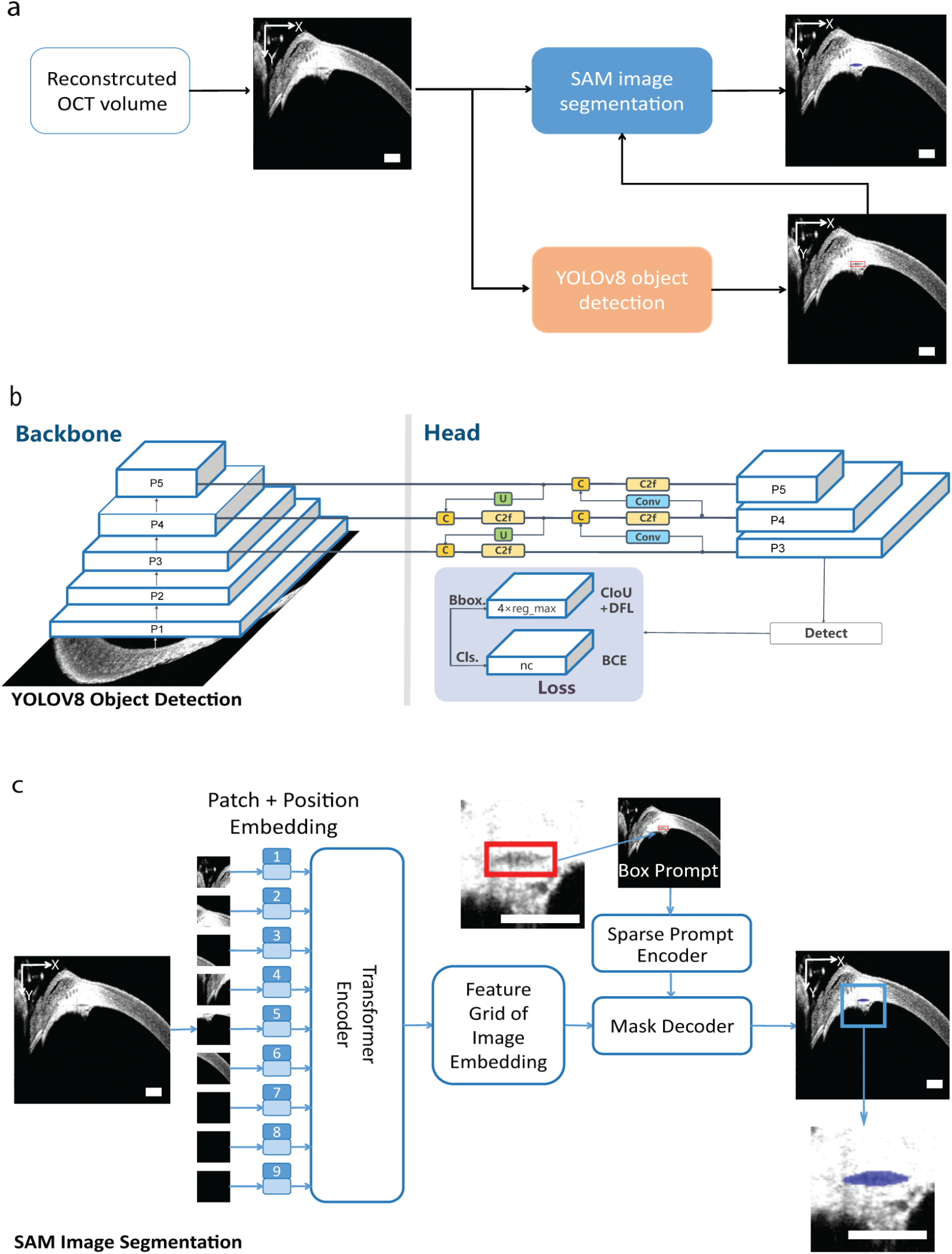
SC-LSS workflow. (a) Overview of the two-stage workflow. Starting from a cross-sectional B-scan from the vis-OCT volume, YOLOv8 was applied to localize a bounding box for SC. The output bounding box was input to SAM for box-promp-based semantic segmentation. SC overlaid in blue. All scale bars are 200 μm. (b) Architecture diagram of the YOLOv8 object detection model. P1-P5 are feature maps at different resolutions extracted by the model. C stands for a convolutional block, Conv stands for a convolutional layer (simple convolution operation), U stands for upsampling block for a feature map and C2f refers to a feature fusion block combining partial convolutions and concatenation. (c) Architecture of SAM image segmentation. B-scan image is divided by Vision Transformer encoder^38^ into 16×16 patches (4096 patches in total). Each patch is added with learnable position embedding and projected into a feature vector. The decoder then combines these feature vectors with the bounding box output from YOLOv8 to generate the final segmentation mask.

During training, we iteratively optimized the YOLOv8 and SAM weights such that the discrepancy between the output segmentation and ground truth is minimized. To enable YOLOv8 and SAM to accurately segment the Schlemm’s canal (SC) in vis-OCT B-scan images, we fine-tuned both models on 324 manually segmentated SC B-scans, allocating 264 images (80%) for training, 30 images (10%) for validation, and 30 images (10%) for testing^37^. We conducted model fine-tuning and inference on a workstation equipped with a 12th Gen Intel® Core™ i7-12700KF CPU (3.61 GHz) and an NVIDIA GeForce RTX 3090 GPU.

### 2.2 YOLOv8 for object detection

We used YOLOv8 to localize the SC position for every vis-OCT B-scan. First, we fine-tuned a pre-trained YOLOv8 model using our vis-OCT SC dataset as a training dataset, where ground truth bounding boxes are generated from the manually segmented SC masks. For each manual segmentation binary mask image, we identified the *x*_*min*_, *z*_*min*_, *x*_*max*_, and *z*_*max*_ coordinates corresponding to the points of the mask, where the *x-*axis corresponds to the horizontal direction in the B-scan and the z-axis to the vertical direction. The bounding box is the rectangle defined by the four corner points (*x*_*min*_, *z*_*min*_), (*x*_*min*_, *z*_*max*_), (*x*_*max*_, *z*_*min*_), and (*x*_*max*_, *z*_*max*_). During training, we compared the bounding boxes generated by YOLOv8 with the ground truth bounding boxes of the training dataset. The YOLOv8 model consists of two major components, the backbone and the head (Fig. 1b). The backbone is the primary feature extractor composed of a convolutional neural network (CNN) that transforms input images into a hierarchy of feature representations. As shown in Fig. 1b, the head of YOLOv8 model is responsible for performing task-specific predictions that map the features extracted from the backbone to bounding boxes through a series of blocks: cross-stage partial with 2-branch feature fusion (C2f)^39^, composite convolutional block (C)^39^, convolution layer (Conv)^39^, and upsampling block (U)^39^.

To enhance the robustness of the fine-tuned YOLOv8 model, we employed a range of data augmentation strategies during training, including random scaling by up to 10% of the image size, translational shifting up to 10% of the image size, horizontal flipping with 50% probability, and brightness changes by plus or minus 40%^39^. We fine-tuned the YOLOv8 model using the Adam optimizer with a learning rate of 0.002, a batch size of 16, and a training schedule of 100 epochs^40^.

For fine-tuning, we minimized a composite multi-task loss consisting of two primary components. The first component is the bounding box regression loss, computed as^39^

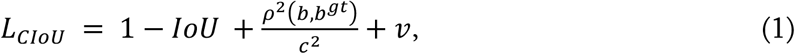

where IoU is the intersection over union; *ρ*^2^(*b, b*^*gt*^) is the Euclidean distance between the center points of the predicted bounding box b outputted from YOLOv8 and the ground truth bounding box *b*^*gt*^ of the manually segmented SC; *c* is the diagonal length of the smallest enclosing box covering both prediction and ground truth bounding box; and *v* is the measure of consistency between ground truth box aspect ratio and predicted box aspect ratio that penalizes the differences in shape. This component penalizes spatial misalignment, aspect ratio differences, and center point discrepancies between predicted and ground truth boxes. We set the weight of this term to 0.05 based on the YOLOv8 default and preliminary experimentation. We selected relative weights for terms to balance the gradient magnitudes of each term in the bounding box regression. The second component is focal loss, measured via binary cross-entropy (BCE) loss as^37^

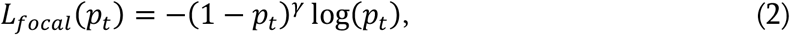

where *p*_*t*_ is the model’s predicted confidence for the correct foreground class and *γ* is a focusing parameter that downweights the easy examples (prevents the model from being overwhelmed by many negatives). The purpose of this component is to distinguish foreground objects from background non-objects, with the weight of this term set as 1.0. After training, we used the fine-tuned YOLOv8 model to generate bounding boxes for SC, which were used as spatial prompts for the downstream semantic segmentation model.

### 2.3 SAM for prompt-based segmentation

For semantic segmentation, the second stage of our pipeline, the input parameter for SAM was a vis-OCT B-scan image and the bounding boxes generated from YOLOv8. We fine-tuned the SAM model using a hybrid prompt strategy that combines bounding boxes and sparse point prompts. Bounding boxes generated by the manually segmented SC masks were reused during training to provide coarse localization guidance, serving as initial spatial constraints to help the model focus on the region likely containing SC. To improve the model’s discriminative capacity, both positive and negative point prompts were incorporated. Positive point prompts are points located inside the SC mask, and negative point prompts are points within the bounding box but outside the SC mask. These prompts assist SAM in determining whether regions containing these points are within or outside SC. We illustrate the steps used by the transformer-based SAM segmentation with a box prompt encoder diagram in Fig. 1c^36^. We split the input image into small non-overlapping patches^38^; then we flattened each patch (i.e., converted the 2D grid of pixel values of the image into a single row of numbers) and added an element-wise positional embedding to record spatial information. The embedded patches pass through the transformer encoder block, which creates a 2D spatial feature grid for use in segmentation. The mask decoder integrates the bounding box determined from YOLOv8 with the image feature grid via transformer layers to output an SC segmentation.

Positive point prompts were generated from each manually segmented mask by computing the center of mass for non-zero pixels of the mask (i.e., the geometric center of the foreground region within the YOLOv8 bounding box). To suppress spurious activations and improve boundary precision, we also introduced negative point prompts at the corners of input bounding boxes. We processed the combined set of bounding boxes and point prompts through the prompt encoder of SAM to produce spatial embeddings for mask decoding^41^. This approach provided the model with spatial cues for foreground (positive) and background (negative) regions, encouraging it to learn more discriminative representations during segmentation mask prediction.

We optimized the SAM model weights by minimizing a composite loss function that included dice loss, binary cross entropy (BCE) loss, total variation (TV), score loss, and false positive penalty. We defined dice loss as^42^

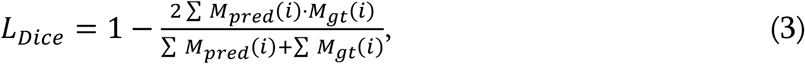

where *M*_*pred*_ is the SAM predicted mask and *M*_*gt*_ is the ground truth mask. Overall, dice loss increases with decreased overlap between segmentations. BCE loss is defined as^37^

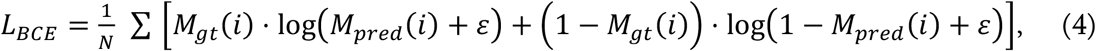

where *N* is the total number of pixels and *ε* is a small constant to ensure the logarithm input is greater than zero. The BCE loss term provides information about pixel-wise probability calibration. We defined TV loss^43^, used to promote spatial smoothness, as

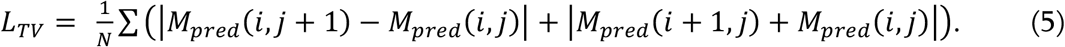

Finally, we have a score loss term defined as^36^

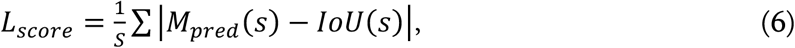

where *S* is the total number of samples; *IoU* is the measured intersection over union for sample s; and *M*_*pred*_ (*s*) is the model-predicted *IoU* score. The total loss L is defined by

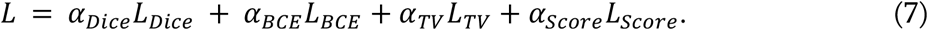

The weights (*α*_*Dice*_, *α*_*BCE*_, *α*_*TV*_, *α*_*Score*_) for all four terms are determined through theoretical normalization. In multi-term loss functions, each component often has a different numeric range, and one term may dominate the training signal without careful balancing. To ensure that all loss terms contributed similarly, we tuned their weights such that their scaled losses had roughly comparable magnitudes during training. This strategy enables balanced gradient updates and stable convergence, preventing dominance by high-magnitude terms^44^.

We trained SAM using the Adam optimizer (learning rate = 0.0001, weight decay = 1e-4) and the gradient accumulation strategy, a technique employed when training models with limited GPU memory, for training^45^. We trained the model for 5000 steps.^37^ During fine-tuning, we validated the model every 50 steps by computing the IoU between the predicted and ground truth masks. To monitor for potential overfitting or model drift, we tracked IoU and training loss. If IoU decreased as the training loss decreased, a sign of model overfitting, we adjusted the learning rate and weight decay parameters to achieve more stable fine-tuning performance.

### 2.4 SC volumetric reconstruction

After generating SC masks for every B-scan for each volume, we took the union of all segmentations to generate a volumetric SC mask. Occasionally, YOLOv8 erroneously identified alternative lumens within the anterior segment of the eye as SC, leading to discontinuities in the 3D reconstruction. We identified these incorrectly localized regions and corrected them based on the 3D continuity of SC segments.

Based on the 3D continuity of true SC, we kept 3D connected components with a volume exceeding 30000 μm^3^ for the volumetric SC mask and eliminated smaller components. B-scans corresponding to these eliminated regions were considered to have a potentially incorrectly localized SC and were subject to a second round of segmentation. For each of these B-scans, we found the closest B-scan in a larger connected region exceeding 30000 μm^3^. We defined the closest B-scan as the B-scan in the larger region with the fewest B-scans in between the B-scans in the eliminated region. Subsequently, we performed rigid registration of the B-scan that was found to have a potentially incorrectly localized SC to the closest B-scan in a larger connected region, with translations for the rigid registration determined by maximizing cross-correlation between images. The bounding box of the correctly localized B-scan was translated according to the offset computed by rigid registration and set as the bounding box of SAM for the B-scans requiring a second segmentation attempt.

### 2.5 Evaluation metrics

We used the Dice coefficient to measure overlap between two segmentations, with a value of 1 indicating perfect overlap and 0 indicating no overlap. Given segmentations A and B, the Dice coefficient is defined as

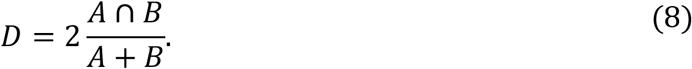

We used the mean surface distance (MSD) to measure the distance between the boundaries of A and B, with an MSD of 0 corresponding to perfectly overlapping boundaries. The directed MSD, the average distance of a point in the boundary of A to its closest point in the boundary of B, is defined as^46^:

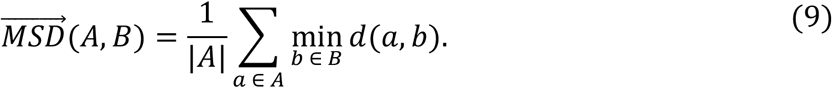

The MSD is the average of the two directed MSDs.

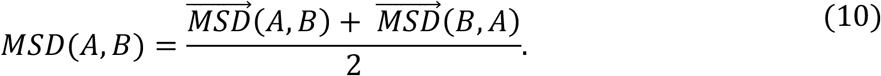

Finally, we used the mean absolute error (MAE) of the SC area and SC height to quantify differences in the morphology of SC. We defined MAE between A and B as the absolute differences in SC area and SC height between A and B. We determined SC height, approximating the distance between the inner and outer walls of SC, by fitting an ellipse to SC and measuring the length of the fitted minor axes^26^.

### 2.6 SC Imaging

*In vivo* imaging of SC was performed using a custom-built robotic anterior segment vis-OCT system as previously described^9, 10, 30^. The lateral FOV for each vis-OCT volume was 2.04 mm, with each volume consisting of 512 A-lines per B-scan and 512 B-scans per volume. We acquired all data using a temporal speckle averaging data acquisition pattern^47^, with an A-line rate of 75 kHz and incident beam power of 1 mW as previously described^9, 10^. Only one eye was imaged per mouse. We anesthetized the mice with an intraperitoneal injection (10mL/kg body weight) of a ketamine/xylazine cocktail (ketamine 11.45mg/L, xylazine 1.7mg/L in saline). For exposure of the limbus, we placed a speculum underneath the eyelids. All imaging protocols were approved by the Northwestern University Institutional Animal Care and Use Committee and complied with the ARVO Statement for the Use of Animals in Vision Research.

### 2.7 Dataset and labeling

We acquired 133 volumes from 15 different adult wild-type C57/BL6J mice aged between 3 and 8 months. In 5 mice, we monometrically varied IOP from 5 mmHg to 20 mmHg in increments of 5 mmHg as previously described^25^. We captured one volume per IOP level per mouse, with a total of 5 volumes acquired at each IOP level. For two mice, we acquired eight volumes around the eye at baseline and eight volumes around the eye 15 minutes after administering 1% pilocarpine, as in our previous work^10^. We used 324 B-scans for training, validating, and testing SC-LSS. In addition to this, a separate independent dataset was used for evaluating the impact of IOP and pilocarpine on SC size. Among the 324 B-scans, 17 were imaged after pilocarpine treatment, 34 at 5 mmHg, 34 at 10 mmHg, 34 at 15 mmHg, 34 at 20 mmHg, and the rest at physiological conditions. For further evaluation of SC-LSS, expert grader 1 segmented SC in a subset of B-scans from the same 133 volumes (n = 606 B-scans, n = 73 at IOP 5 mmHg, n = 75 at IOP 10 mmHg, n = 76 at IOP 15 mmHg, n = 59 at IOP 20, n = 112 before pilocarpine adminstration, and n = 129 B-scans post pilocarpine administration, rest at physiological conditions). In a random subset of 15 of the acquired volumes (2 at IOP 5 mmHg, 2 at IOP 10 mmHg, 1 at IOP 15 mmHg, 1 at IOP 20 mmHg, 2 after pilocarpine administration, and 7 without any perturbation), three expert graders segmented out SC every 30 B-scans (n = 203 B-scans). We generated manual segmentations by importing the vis-OCT volumes into MATLAB’s *Volume Segmenter* application, delineating the SC boundaries with the paintbrush tool within the application, and exporting the segmentations as PNG images. In cases with multiple manual segmentations, the ground truth was determined with majority voting among all manual segmentations. In cases with only a single manual segmentation, the ground truth was the same as the single manual segmentation.

### 2.8 Statistics

When comparing two values, we performed a paired two-tailed T-test to assess statistical significance unless otherwise specified. When comparing values at baseline and after pilocarpine administration, we performed an unpaired two-tailed T-test to assess statistical significance. When comparing differences in SC area, Dice coefficient, and MSD at various IOPs, we performed a one-way ANOVA analysis. Tukey correction was applied to account for multiple comparisons. When assessing the relationship between SC size, Dice, MSD, and MAE of SC area and height, we performed a Pearson’s correlation analysis. All statistical tests were performed using GraphPad Prism 10.4.0, and all reported values are presented as the mean ± standard deviation.

## 3 Results and discussion

### 3.1 Comparison with human graders

We compared the SC segmentation of SC-LSS with those of two expert graders. SC is a hypo-reflective lumen carrying aqueous humor immediately lateral to the iridocorneal angle. Figure 2 shows the segmentation of SC for two example cross-sectional B-scans. In the first B-scan, we observed two hypo-reflective vascular lumens lateral to the SC in the sclera and a tear meniscus superior to the SC (Fig. 2a). Without the localization step of SC-LSS, these structures could be confused with the SC and the iris, based on their reflectivity and shape. In this example, the expert graders and SC-LSS produced similar segmentation results (Figs. 2b-2c). In the second example, we observed a specular reflection medial to SC in the cornea (Fig. 2d). The specular reflection did not impact the segmentation, as both expert graders and SC-LSS produced similar segmentations (Fig. 2e-f).

**Figure 2.**
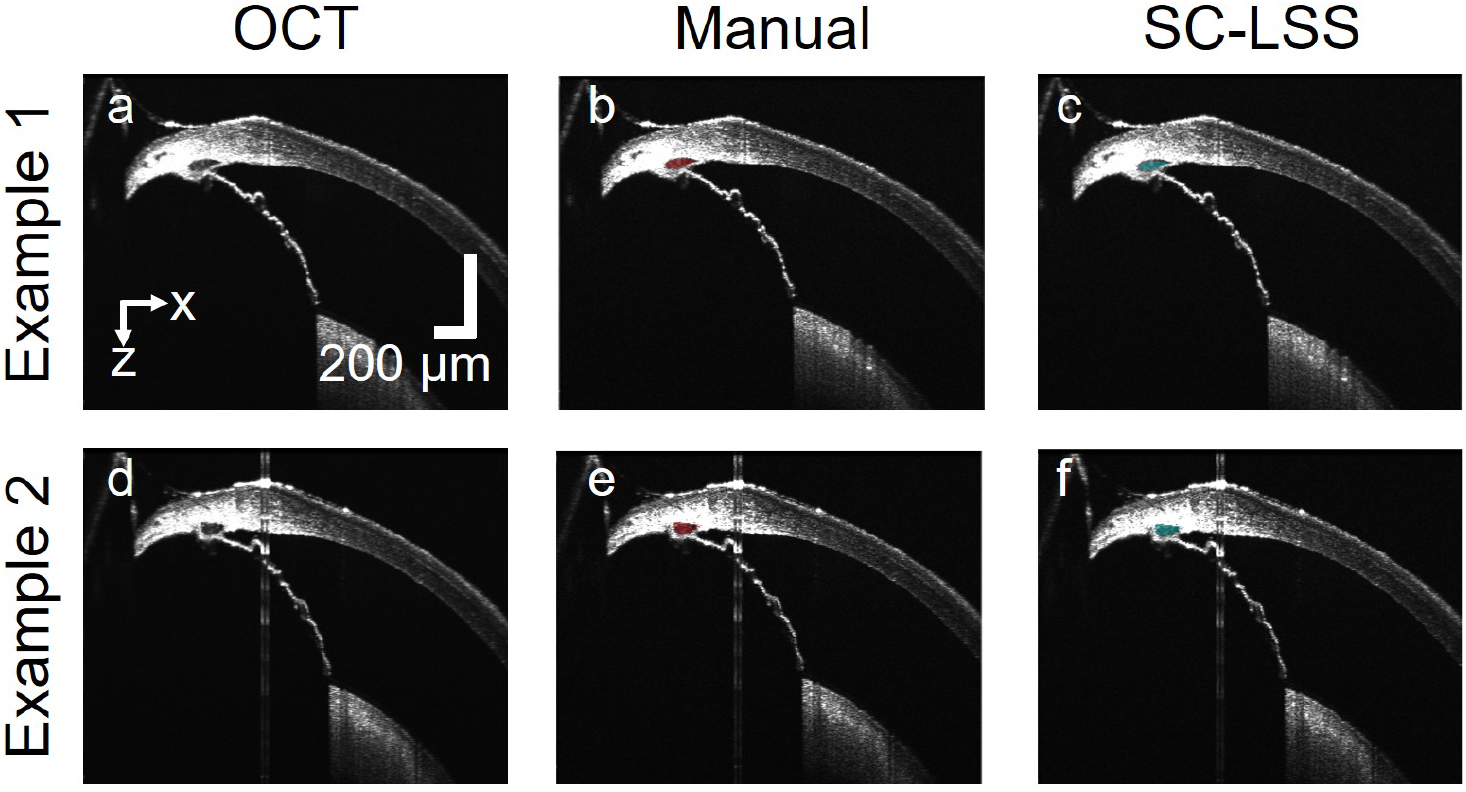
Comparison between expert grader and SC-LSS segmentations. (a) and (d) Vis-OCT cross-sectional B-scan images; (b) and (e) manual segmentation of SC overlaid in red; (c) and (f) SC-LSS segmentation of SC overlaid in cyan. We note different lengths of scale bars in the x and z axes, as the length of the SC in the x direction is larger than in the z direction.

Quantitatively, the difference between SC-LSS and expert grader segmentation results was comparable to the inter-observer variability. We measured a Dice coefficient of 0.70±0.20, a mean surface distance of 6.0±7.3 µm, and an MAE of 589±763 µm^2^ in cross-sectional SC area and an MAE of 4.7±5.6 µm in SC height between the ground truth and SC-LSS (n = 203 B-scans). Furthermore, we measured an average Dice coefficient of 0.73±0.18 and an average mean surface distance of 6.6±11.1 µm between the three graders (n = 203 B-scans). Neither the Dice coefficient (p = 0.10) nor the mean surface distance was statistically significant between the two groups (p = 0.48). Overall, the computation time of SC-LSS was 0.27 seconds per B-scan and 70 seconds per volume.

### 3.2 Reconstruction of 3D Volumes

Next, we assessed the capacity of SC-LSS to create 3D reconstructions of SC. Fig. 3a shows a 3D reconstruction of SC generated by SC-LSS, with the 3D segmentation continuous as expected. For segmented B-scans within all volumes, 91.7±7.1% of segmented B-scans had at least 50% overlap in the segmented region with neighboring B-scans (n = 133 volumes). The average overlap, as measured by Dice coefficient, between segmented B-scans was 83.1±6.6% (n = 133 volumes). Furthermore, 73.7±9.7% of segmented SC had a greater percentage overlap with SC in neighboring B-scans than with SC in B-scans separated by one intermediary B-scan (n = 133 volumes).

**Figure 3.**
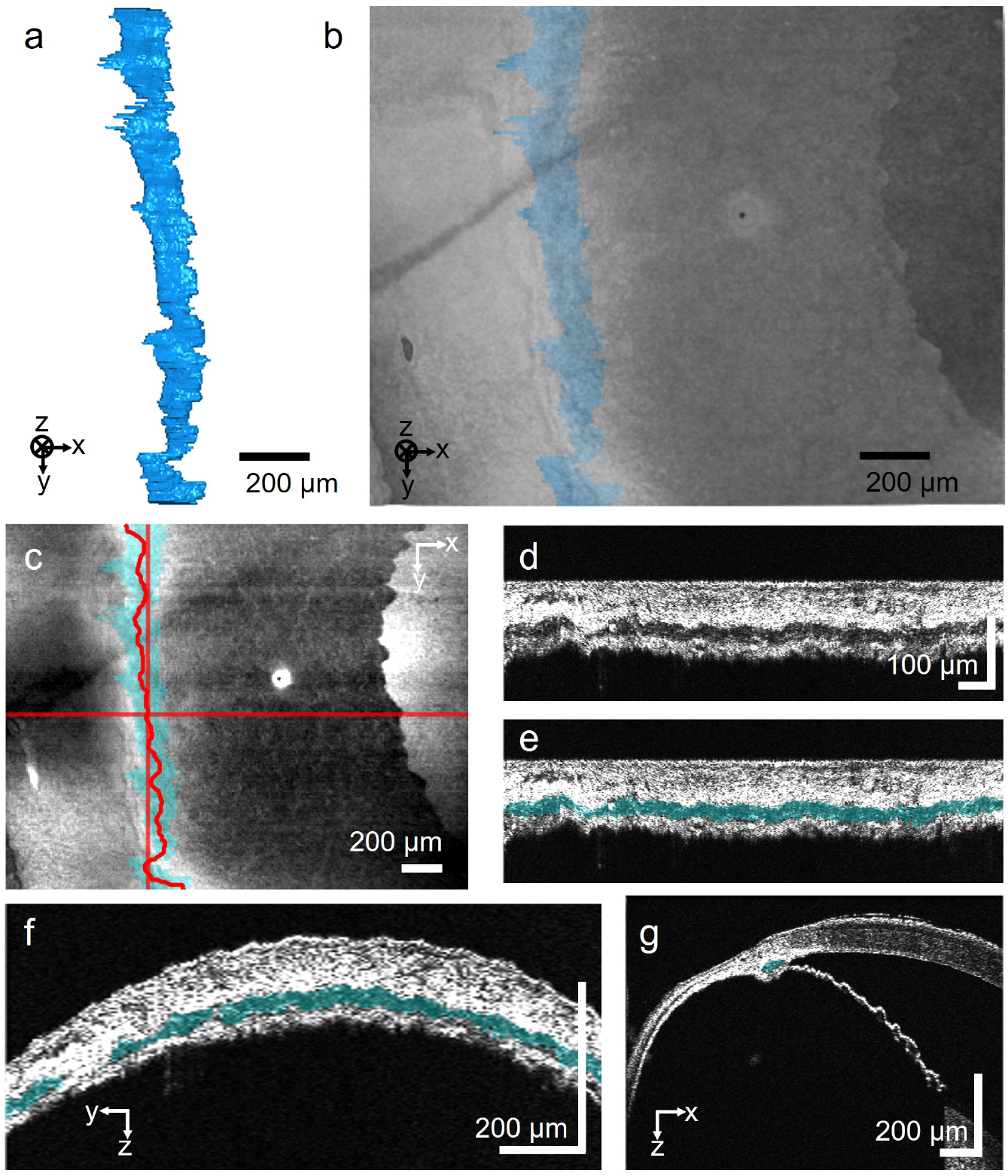
Three-dimensional volumetric reconstruction of SC. (a) 3D volumetric representation of SC generated by SC-LSS; (b) Segmented SC inside 3D volumetric reconstruction of OCT data. (c) En-face projection of the OCT volume with the position of SC dictated by the blue overlay. (d) Cross-section through the center of SC following the red curve, which is the skeleton of the SC segmentation in (c). (e) Segmentation of SC overlaid in blue. (f) Cross-sectional B-scans corresponding to the vertical red line in (c). (g) Cross-sectional B-scans corresponding to the horizontal red line in (c).

Figure 3c shows the *en face* projection of the same OCT volume with the position of SC overlaid in blue. For this volume, the 3D reconstruction of SC remained continuous. We viewed a digitally resampled flattened cross-section of a plane passing through the center of SC (Fig. 3d)^9^ and overlaid the segmentation of SC through that plane in blue (Fig. 3e). Even though data from the resampled cross-section came from over three hundred different B-scans, each segmented individually, the segmentation of SC in the resampled cross-section conformed to the boundaries of SC. The bottom of the segmentation is the inner wall of SC, and the top of the segmentation is the outer wall of SC. Fig. 3f shows a B-scan along the direction of the slow scanning mirror of the OCT volume, with segmentation of SC following the visible boundaries of SC despite noticeable motion artifacts. As expected, qualitative observation of SC segmentation along each B-scan within the volume aligned with observer expectations (Fig. 3g).

### 3.3 Evaluation of change in SC morphology subjected to IOP modulation

We further evaluated changes in SC morphology in response to IOP modulation detected by SC-LSS. We show cross-sectional B-scans of the mouse limbus at IOP ranging from 5-20 mmHg (Figs. 4a-4d) with the segmentations generated by SC-LSS overlaid in blue (Figs. 4e-4h). We observed that the segmented SC size decreases with increasing IOP, consistent with the literature^25, 26^. Although SC morphology varies significantly across different IOPs, SC-LSS-generated segmentations consistently conformed to SC boundaries at all IOPs. SC-LSS measured the average SC area of 2235±1185 μm^2^ and SC height of 20.3 ± 5.4 μm at 5 mmHg (n = 73 B-scans). Compared to the SC area at 5 mmHg, SC-LSS found that SC area was 1656±873 μm^2^ at 10 mmHg, 25.9% smaller (n = 75 B-scans; P = 0.0002 compared to 5 mmHg) than at 5 mmHg. It found that SC height was 17.3±4.9 μm at 10 mmHg, 14.4 % smaller (P = 0.002 compared to 5 mmHg) than at 5 mmHg. At 15 mmHg, SC-LSS measured an SC area of 1152±688 μm^2^, 48.4% smaller (n = 76 B-scans; P < 0.0001 compared to 5 mmHg) than at 5 mmHg. It found that SC height was 14.7±4.5 μm at 15 mmHg, 27.8 % smaller (P < 0.0001 compared to 5 mmHg) than at 5 mmHg. At 20 mmHg, SC-LSS measured an SC area of 1082 ± 664 μm^2^, 51.5% smaller (n = 59 B-scans; P < 0.0001 compared to 5 mmHg) than at 5 mmHg (Fig. 4i). It found that SC height was 14.4 ± 4.6 μm at 20 mmHg, 28.8 % smaller (P < 0.0001 compared to 5 mmHg) than at 5 mmHg.

**Figure 4.**
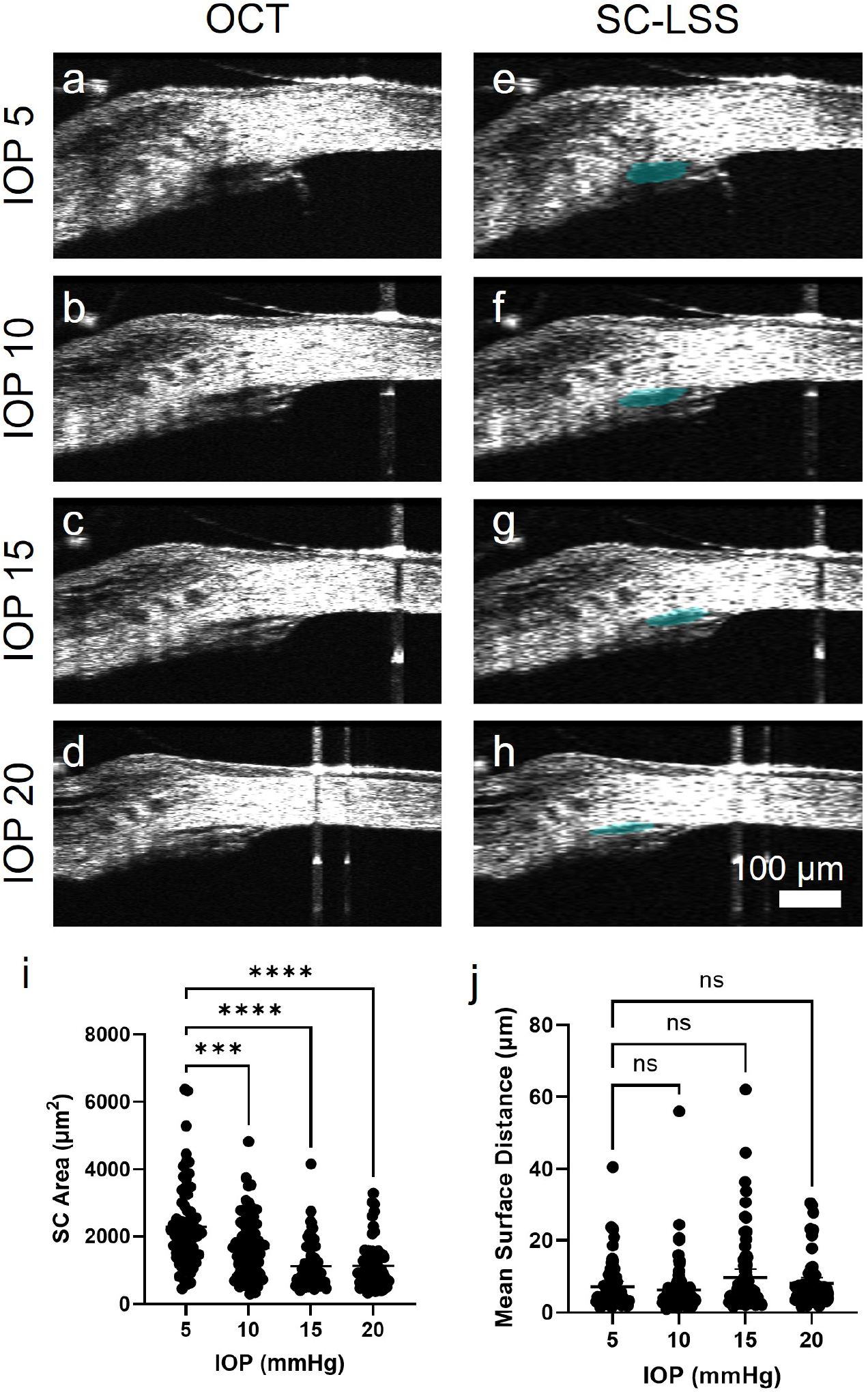
SC-LSS is sensitive to IOP-induced changes in SC morphology. (a-d) B-scan taken at IOP = 5, 10, 15, and 20 mmHg with (e-h) SC-LSS segmented SC overlaid in blue; (i) SC area reduces with elevated IOP; (j) MSD is relatively stable with elevated IOP. ns > 0.05, * < 0.05, ** < 0.01, *** < 0.001, **** < 0.0001.

We examined the relationship between segmentation accuracy, IOP, and SC size. We calculated the Dice coefficient between SC-LSS and the expert grader 1 as 0.68±0.18 at 5 mmHg (n = 73 B-scans), 0.64±0.18 at 10 mmHg (n = 75 B-scans, P = 0.68 compared to 5 mmHg), 0.52±0.22 at 15 mmHg (n = 76 B-scans, P < 0.0001 compared to 5 mmHg), and 0.45±0.21 at 20 mmHg (n = 59 B-scans, P < 0.0001 compared to 5 mmHg). The MSD between SC-LSS and manual grader 1 was 7.2±6.6 μm at 5 mmHg, 6.3±7.3 μm at 10 mmHg (P = 0.90 compared to 5 mmHg), 9.7±10.7 μm at 15 mmHg (P = 0.24 compared to 5 mmHg), and 8.1±6.8 μm at 20 mmHg (P = 0.93 compared to 5 mmHg; Fig. 4j). One-way ANOVA revealed a statistically significant reduction in the Dice coefficient (P < 0.0001) between SC-LSS and expert graders with increasing IOP. We noted a trend of increasing MSD between SC-LSS and manual graders with increasing IOP, but it was not statistically significant (P = 0.068). Indeed, we found a positive Pearson correlation (R = 0.45, P < 0.0001) between Dice coefficient and SC area; however, there was no statistically significant correlation between MSD and SC area (R = −0.04, P = 0.55).

In aggregate, the decreasing Dice coefficient with decreasing SC size suggests that SC-LSS might perform more poorly when SC size is smaller. However, the stable relationship between MSD, a measure of how close the boundaries of segmentation generated by SC-LSS and the ground truth, and IOP and SC area suggests that the ability of SG-LSS to identify the boundaries of SC is not significantly impacted by SC size. Reconciling these observations, misidentifying the boundaries of SC by a fixed distance will have a greater impact on Dice when SC is smaller than when it is larger.

### 3.4 Evaluation of change in SC morphology with pilocarpine administration

Moreover, we evaluated changes in SC morphology in response to pilocarpine administration detected by SC-LSS. We captured baseline OCT B-scans with SC (Figs. 5a-5b) and subsequently captured OCT B-scans of the same regions after 1% pilocarpine administration (Figs. 5c-5d). The average Dice coefficient and MSD were 0.70 ± 0.22 and 7.6 ± 11.4 μm at baseline (n = 112 B-scans), and 0.73 ± 0.20 and 6.1 ± 7.7 μm after pilocarpine administration (n = 129 B-scans), with no statistical significance found in Dice coefficient (P = 0.29) and MSD (P = 0.22) between the two groups (Fig. 5e-f). The MAE in SC area was 644 ± 478 μm^2^ at baseline, and 700 ± 829 μm^2^ after pilocarpine administration, with no statistical significance found (P = 0.53) between the two groups. The MAE in SC height was 3.3 ± 2.6 μm at baseline, and 3.7 ± 6.0 μm after pilocarpine administration, with no statistical significance found (P = 0.44) between the two groups. SC-LSS measured a mean SC area of 2824 ± 1556 μm^2^ and SC height of 19.6 ± 6.8 μm at baseline and a SC area of 3391 ± 1821 μm^2^ and SC height of 22.8 ± 8.6 μm after pilocarpine administration. Consistent with the literature results^28, 48^, SC-LSS found that pilocarpine increased SC size (P = 0.011; Fig. 5g) and SC height (P = 0.002).

**Figure 5.**
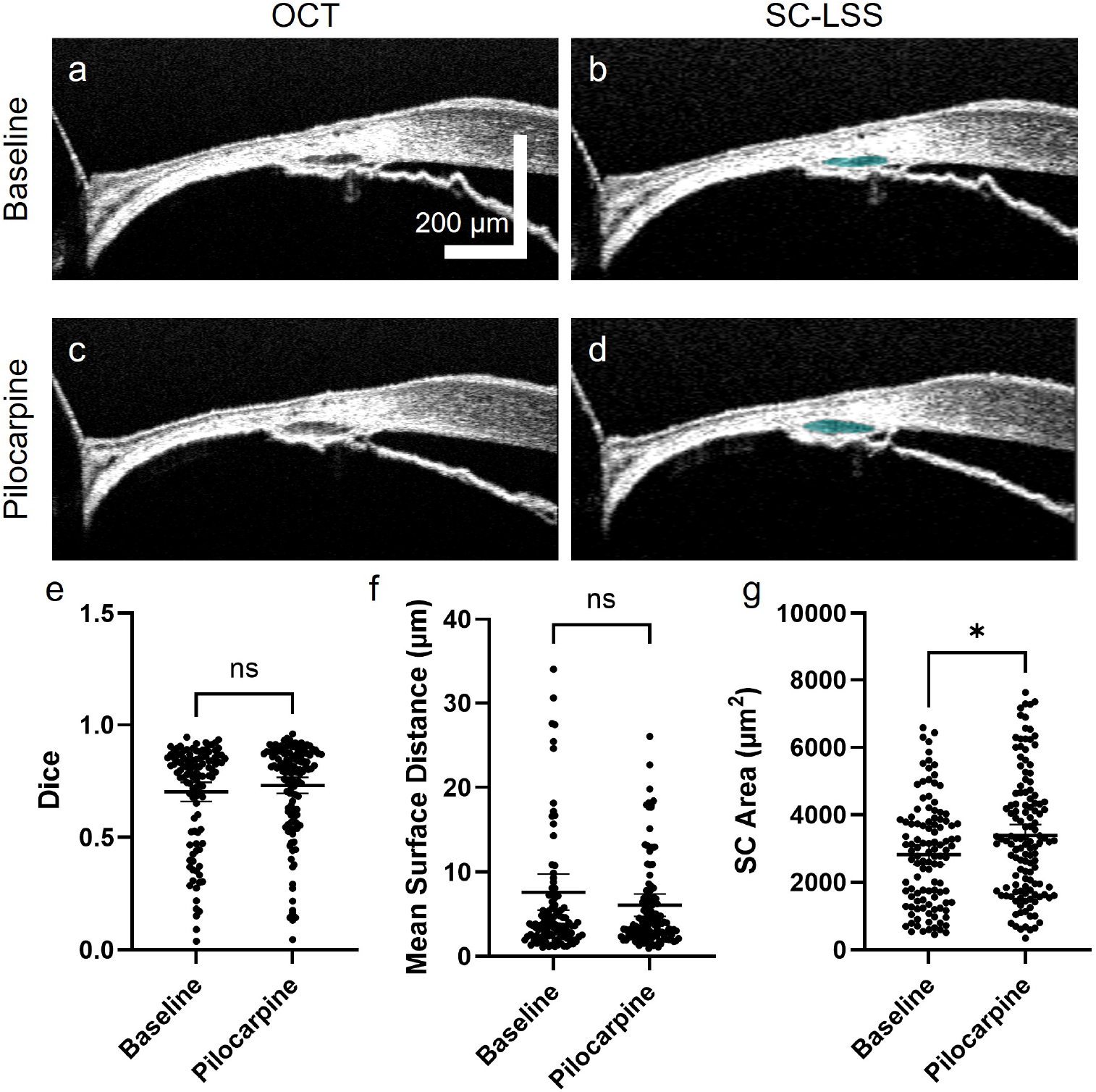
SC-LSS is sensitive to pilocarpine-induced SC morphology changes. (a) Vis-OCT B-scan before pilocarpine administration with (b) SC-LSS segmented SC overlaid in blue; (c) B-scan in identical position after pilocarpine administration with (d) SC-LSS segmented SC overlaid in blue; (e) Impact of pilocarpine on segmentation accuracy as measured by Dice coefficient; (f) MSD; and (g) the SC area. ns > 0.05, * < 0.05.

### 3.5 Discussion

Overall, we observed that SC-LSS produced accurate SC segmentations for vis-OCT volumes. With over 500 B-scans per acquired vis-OCT volume, manual segmentation of the entire volume may take days. However, SC-LSS can segment the full volume in about a minute without requiring manual intervention. Investigating the trabecular outflow pathway based on volumetric SC segmentation instead of a few representative cross-sectional SC segmentations will enable more accurate studies regarding the known segmental differences in its anatomy and the potential segmental differences in its functions. This feature has the potential to guide patient-specific decisions in glaucoma surgical management^49–54^. However, we do note several limitations of our study. First, all vis-OCT volumes used in this study were obtained from a single institution using one vis-OCT system. Second, the training and testing data used in this study were obtained from 16 eyes, which is a relatively small number, and can potentially limit the generalizability of the SC-LSS. Thus, SC-LSS would benefit from a more diverse set of training data. As vis-OCT devices become increasingly widely available, access to a broader range of data from various institutions and experimental conditions will increase, thereby enhancing the external validity of our model. Acquiring data from a wide range of experimental conditions is important because various factors influence IOP and SC size *in vivo*, including the type of anesthesia, head positioning, time of day, and handling of the mice^25, 55–57^. Under alternative experimental conditions, where measured SC might be smaller, the model would benefit from additional training. We note that the SC size measured in this work solely depended on expert graders manually segmenting the SCs from *in vivo* vis-OCT B-scan images, which may be different from that measured from *ex vivo* images, such as histology and fluorescence microscopy. Further studies are required to thoroughly investigate the size of SC *in vivo* and to establish the gold-standard values of the SC size in mice under different conditions.

SC-LSS measured changes in SC size with pilocarpine administration and IOP modulation. SC-LSS measurement of 20.1% increase in SC size after pilocarpine administration, which is on the lower end of published values in humans and mice. Other studies have reported a 26.8%^10^, 131.6%^28^, 21.0%^48^, and 24.7%^58^ increase in size. Additionally, SC-LSS measured a 48.4% decrease in SC size when IOP was increased from 5 mmHg to 15 mmHg, which is less than the 72% decrease in SC size noted with an elevation of IOP of 10 mmHg in a previous study^25^. Currently, SC-LSS might overestimate the SC area at higher IOP, likely because there were far less training data at higher IOP values than at the physiological conditions. We expect increased training with the higher IOP or a separate model for higher IOP would improve results. Finally, SC-LSS compares favorably to other fully automated SC segmentation methods. Choy et al. found a similar Dice coefficient between a machine learning-based method and inter-observer variability, but their study involved a much smaller FOV and consisted of a single cross-sectional image averaged 200 times per eye, rather than the full volume^34^. Zambrano et al. developed a machine learning-based automatic segmentation methodology for humans, but their current Dice is around 0.4, which is significantly lower than our value of 0.7, albeit on much different samples^59^.

In the future, we plan to expand our SC-LSS to include segment alternative elements of the trabecular outflow pathways, such as the TM, collector channels, aqueous veins, and episcleral veins^60–62^. Additionally, we plan to expand the application of our model to volumes acquired from conventional near-infrared OCTs, which currently have more widespread usage than vis-OCT. Ultimately, we aim to adapt our segmentation methodology for SC imaging in humans, with the long-term goal of utilizing SC morphology to inform clinical decision-making.

## 4. Conclusions

In summary, we developed SC-LSS for automatic segmentation of volumetric SCs for vis-OCT. SC-LSS consists of two processes: it first identifies the region with SC and then identifies the boundaries of SC within the area. Finally, it merges the segmentation of each cross-section to form a volumetric segmentation of the SC. Using the continuity of SC, outlier segmentations were identified, the bounding box of the region containing SC was corrected, and a new cross-sectional segmentation was generated. Overall, the performance of SC-LSS in SC segmentation was comparable to that of expert human graders, and it was able to identify expected SC morphology changes in response to IOP modulation and pilocarpine administration. SC-LSS reduces the time-intensive step of SC segmentation, thereby accelerating preclinical studies that focus on the trabecular outflow pathways with a focus on advancing our understanding of pathologies impacting the trabecular outflow pathways, such as glaucoma.

## Acknowledgement

The authors sincerely acknowledge the generous support from NIH grants R01EY033813, R01EY034740, R01EY030501, R01EY03050, U01EY033001, T32GM142604, F30EY034033, Research to Prevent Blindness/David L. Epstein Career Advancement Award in Glaucoma Research sponsored by Alcon, American Glaucoma Society Mid-Career Physician Research Grant, and an unrestricted grant from Research to Prevent Blindness (New York, NY).

## Author Information

### Authors

Raymond Fang: Department of Biomedical Engineering, Northwestern University-2145 Sheridan Road, Evanston, Illinois, USA 60208.

Department of Ophthalmology, Northwestern Feinberg School of Medicine – 259 E Erie St, Chicago, Illinois, USA. 60611.

Fengxuanshan Xu: Department of Biomedical Engineering, Northwestern University-2145 Sheridan Road, Evanston, Illinois, USA 60208

Zihang Yan: Department of Biomedical Engineering, Northwestern University-2145 Sheridan Road, Evanston, Illinois, USA 60208

Cheng Sun: Department of Mechanical Engineering, Northwestern University-2145 Sheridan Road, Evanston, Illinois, USA 60208

Tsutomu Kume: Department of Medicine, Northwestern Feinberg School of Medicine– 259 E Erie St, Chicago, Illinois, USA 60611

Alex S. Huang: Department of Ophthalmology, University of California, San Diego-9415 Campus Point Dr, La Jolla, CA, USA 92093

### Author Contributions

The manuscript was written through the contributions of all the authors. All authors have approved the final version of the manuscript. RF, FX, and ZY acquired and segmented the experimental data. FX, RF, and HFZ developed the methodology. TK, CS, and AH contributed to data preparation and machine learning training. ‡ RF and FX contributed equally.

### Notes

HFZ and CS have financial interests in Opticent Health, which is commercializing Vis-OCT; however, Opticent Health did not fund this work. AH is a consultant for Allergan, Amydis, Celanese, Equinox, Glaukos, QLARIS, Santen, Spinogenix, and Topcon and has financial support from Diagnosys, Glaukos, and Heidelberg Engineering.

## Abbreviations

SC: Schlemm’s Canal
SC-LSS: Schlemm’s Canal Attention-Gated-Transformer
TM: Trabecular Meshwork
OCT: Optical Coherence Tomography
vis-OCT: Visible Light-Optical Coherence Tomography
FOV: Field-of-view
MAE: Mean Absolute Error
MSD: Mean Surface Distance
IoU: Intersection Over Union.

